# Epigenome-wide analysis uncovers a blood-based DNA methylation biomarker of lifetime cannabis use

**DOI:** 10.1101/620641

**Authors:** Christina A. Markunas, Dana B. Hancock, Zongli Xu, Bryan C. Quach, Dale P. Sandler, Eric O. Johnson, Jack A. Taylor

## Abstract

Cannabis use is highly prevalent and is associated with adverse and beneficial effects. To better understand the full spectrum of health consequences, biomarkers that accurately classify cannabis use are needed. DNA methylation (DNAm) is an excellent candidate, yet no blood-based epigenome-wide association studies (EWAS) in humans exist. We conducted an EWAS of lifetime cannabis use (ever vs. never) using blood-based DNAm data from a case-cohort study within Sister Study, a prospective cohort of women at risk of developing breast cancer (Discovery N=1,730 [855 ever users]; Replication N=853 [392 ever users]). We identified and replicated an association with lifetime cannabis use at cg15973234 (*CEMIP*): combined P=3.3×10^−8^. We found no overlap between published blood-based *cis*-meQTLs of cg15973234 and reported lifetime cannabis use-associated SNPs (P<0.05), suggesting that the observed DNAm difference was driven by cannabis exposure. We also developed a multi-CpG classifier of lifetime cannabis use using penalized regression of top EWAS CpGs. The resulting 49-CpG classifier produced an area under the curve (AUC)=0.74 (95% confidence interval [0.72, 0.76], P=2.00×10^−5^) in the discovery sample and AUC=0.54 ([0.51, 0.57], P=4.64×10^−2^) in the replication sample. Our EWAS findings provide evidence that blood-based DNAm is associated with lifetime cannabis use.

## INTRODUCTION

Cannabis use is highly prevalent with 45% of Americans aged 12 or older reporting lifetime use (defined here as ever-use of cannabis), and 15% reporting use in the past year^1^. These numbers are expected to grow due to increasing legalization of both medical and recreational cannabis use. As of 2019, 33 US States have legalized cannabis for medical purposes, 11 of which have also legalized recreational use (Accessed 08/14/19^2^). Adverse health effects of cannabis use have been reported for short-term use (e.g., impaired cognitive and motor function, altered judgement, paranoia, and psychosis)^3^, long-term or heavy use (e.g., increased risk of cannabis use disorder, altered brain development, cognitive impairment, chronic bronchitis)^3^, as well as lifetime (ever) use (e.g., psychotic disorder)^4^. In contrast, evidence exists supporting therapeutic benefits for various clinical conditions (e.g., glaucoma, acquired immune deficiency syndrome, nausea, chronic pain, inflammation)^3^. Efforts to better understand the full spectrum of cannabis-related health consequences have been hindered as a result of under-reporting (e.g., due to social stigma associated with use) and the absence of available biomarkers that can accurately quantify cannabis usage patterns.

Currently available biomarkers of cannabis exposure, such as urinary metabolites, suffer limitations. These existing biomarkers have limited windows for detection, are largely restricted to acute exposures (e.g., ranging from 3 to >30 days, depending on the frequency of cannabis use^5^), and are unable to quantify cumulative exposure, which may prove to be a better indicator of subsequent health outcomes^5^.

Epigenetic modifications represent promising candidates for biomarker research. DNA methylation (DNAm), the most commonly studied form of epigenetic modification, is defined by the presence of a methyl group (-CH_3_) most often at the carbon-5 position of a cytosine nucleotide in the context of CpG sites (adjacent cytosine and guanine nucleotides linked by a phosphate group). DNAm can be influenced by genetic factors, disease, environmental exposures, and lifestyle, and can vary across developmental stages of life (e.g., infancy, childhood, and adulthood), tissues, and cell types. An important feature of exposure-related DNAm changes is that they can either be persistent (i.e., stable changes) or reversible (i.e., return to prior state) once the exposure is no longer present. This combination of both persistent and reversible changes has value for biomarker development and has been observed, for example, in relation to tobacco smoking where only a subset of DNAm changes identified between current vs. never smokers are also found between former vs. never smokers^6,7^.

Research geared towards understanding epigenetic changes related to cannabinoids, which encompass endocannabinoids (endogenous ligands), natural cannabinoids (derived from cannabis and includes Δ^9^-tetrahydrocannabinol [THC] and cannabidiol), and synthetic cannabinoids, is growing. However, studies have been largely based on animal studies and human candidate gene studies^8,9^, with the only epigenome-wide study of cannabis use being performed in human sperm^10^. While not specific to cannabis use, there has also been one blood-based genome-wide longitudinal DNAm study in humans which identified early life DNAm changes associated with later adolescent substance use (latent factor combining tobacco, cannabis and alcohol use)^11^. To begin to address this gap in the field, we report the first blood-based EWAS of lifetime cannabis (ever vs. never) use, conducted using discovery (N = 1,730) and replication (N = 853) samples. We extended our EWAS findings by using genetic information to help distinguish genetically-vs. exposure-driven effects. We further leveraged our EWAS results to develop the first multi-CpG classifier of lifetime cannabis (ever vs. never) use.

## METHODS

### Study population

The Sister Study is a prospective cohort of 50,884 women ascertained across the US between 2003–2009 that was designed to examine environmental and genetic risk factors of breast cancer^12^. Women, aged 35–75 years old, were enrolled if they had a sister diagnosed with breast cancer and no personal history of breast cancer at baseline. Whole blood samples were collected, along with data from questionnaires and interviews covering demographics, lifestyle, family and medical history, and environmental exposures.

Since the study’s inception, a number of sub-studies have been designed to address specific hypotheses related to women’s health. For the current study, we used existing DNAm array data from a case-cohort study of 2,878 non-Hispanic white women designed to identify blood-based DNAm changes associated with incident breast cancer, as described previously^13,14^. Briefly, the case-cohort study included a random sample of 1,336 women from the cohort and 1,542 additional women who later developed *in situ* or invasive breast cancer during follow-up (between enrollment and sampling of the case-cohort in 2015). Out of the 2,878 women, 1,616 women (1,542 + 46 cases from the random sample) were diagnosed with breast cancer during follow-up and 1,262 women (1,336 – 46 cases from the random sample) remained clinically breast cancer free (see Supplementary Fig. S1 for study design). All women included in this study were clinically breast cancer free at the time of the blood draw used for DNAm data generation.

Informed written consent was obtained from all Sister Study participants. The Sister Study was approved by the Institutional Review Boards (IRBs) of the National Institute of Environmental Health Sciences (NIEHS), National Institutes of Health, and the Copernicus Group (http://www.cgirb.com/irb-services/). The current study was approved by the IRB at RTI International. All research was performed in accordance with relevant guidelines and regulations.

### Cannabis assessment

Lifetime cannabis use was determined by response to the question, “*Have you <u>ever</u> smoked marijuana?*”, asked along with questions about tobacco smoking during the baseline computer-assisted telephone interview. Basing our primary analysis on ever-use facilitated our evaluation of genetic data using the largest genome-wide association study (GWAS) of cannabis use reported to date (N = 184,765) which focused on ever-use only^15^. Age of initiation (*“How old were you the first time you smoked marijuana?”*), duration of use (*“In total, how many years did you smoke marijuana?”*), and frequency of use (*“During the years that you smoked marijuana, on average how often did you smoke it?”*) were also assessed in the Sister Study and were considered for secondary analyses in the current study to characterize top EWAS findings. No information was available on time since last use to classify ever cannabis users as current vs. former users. Sister Study questionnaires can be accessed at https://www.sisterstudystars.org/.

### DNAm data, quality control (QC), and pre-processing

Blood sample collection and DNA extraction have been described previously^13,14^. A total of 2,878 blood samples were assayed using the Illumina HumanMethylation450 array which covers >450,000 CpG sites targeting promoters, CpG islands, 5’ and 3’ untranslated regions, the major histocompatibility complex, and some enhancer regions^16^.

Data quality assessment and pre-processing were conducted using the R package, *ENmix*^17^. A series of diagnostic plots were generated to detect problematic samples, arrays, and laboratory plates. The ENmix pipeline was implemented for data pre-processing and included the following steps: background correction using the ENmix method^17^; correction of fluorescent dye-bias using the RELIC method^18^; inter-array quantile normalization; and correction of probe type bias using the RCP (regression on correlated CpGs) method^19^. Surrogate variables (SVs) of the array control probes were generated to adjust for technical artifacts. To control for cellular heterogeneity, blood cell type proportions (CD8T cells, CD4T cells, natural killer cells [NK], B cell, monocytes [Mono], and granulocytes [Gran]) were estimated following the Houseman method^20^. β-values, corresponding to the ratio of methylated intensities relative to the total intensity, were calculated to represent DNAm levels at each CpG.

Both sample- and probe-level exclusions were applied. In total, 295 samples were excluded due to poor bisulfite conversion efficiency (average bisulfite intensity <4000), outlier based on QC diagnostic plots (e.g., DNAm β-value distribution), low call rate (> 5% low quality data [Illumina detection P>1×10^−6^, number of beads <3, or values outside 3*Interquartile range (IQR)]), related individuals (one sister from each pair was selected at random for exclusion), missing phenotype data, or date of breast cancer diagnosis preceding blood draw. A total of 67,564 probes were excluded: 16,100 probes with > 5% low quality data; 14,522 probes with a common SNP (minor allele frequency >0.01) at the single base extension site or CpG site; 26,799 cross-reactive probes mapping to multiple genomic locations^21^; 10,143 probes mapping to sex chromosomes. In addition, extreme DNAm β-value outliers were removed (outside 3*IQR); missing values were imputed using K-nearest neighbor. Following exclusions, the final analysis dataset included 2,583 women (Supplementary Fig. S1) and 417,948 CpGs.

### EWAS analysis

The Sister Study breast cancer case-cohort sample was randomly divided into discovery (2/3 of the overall sample: N = 1,730 [855 ever users]) and replication (1/3 of the overall sample: N = 853 [392 ever users]). Characteristics of the case-cohort sample used for analysis are provided in Table 1 and Supplementary Table S1.

**Table 1.**
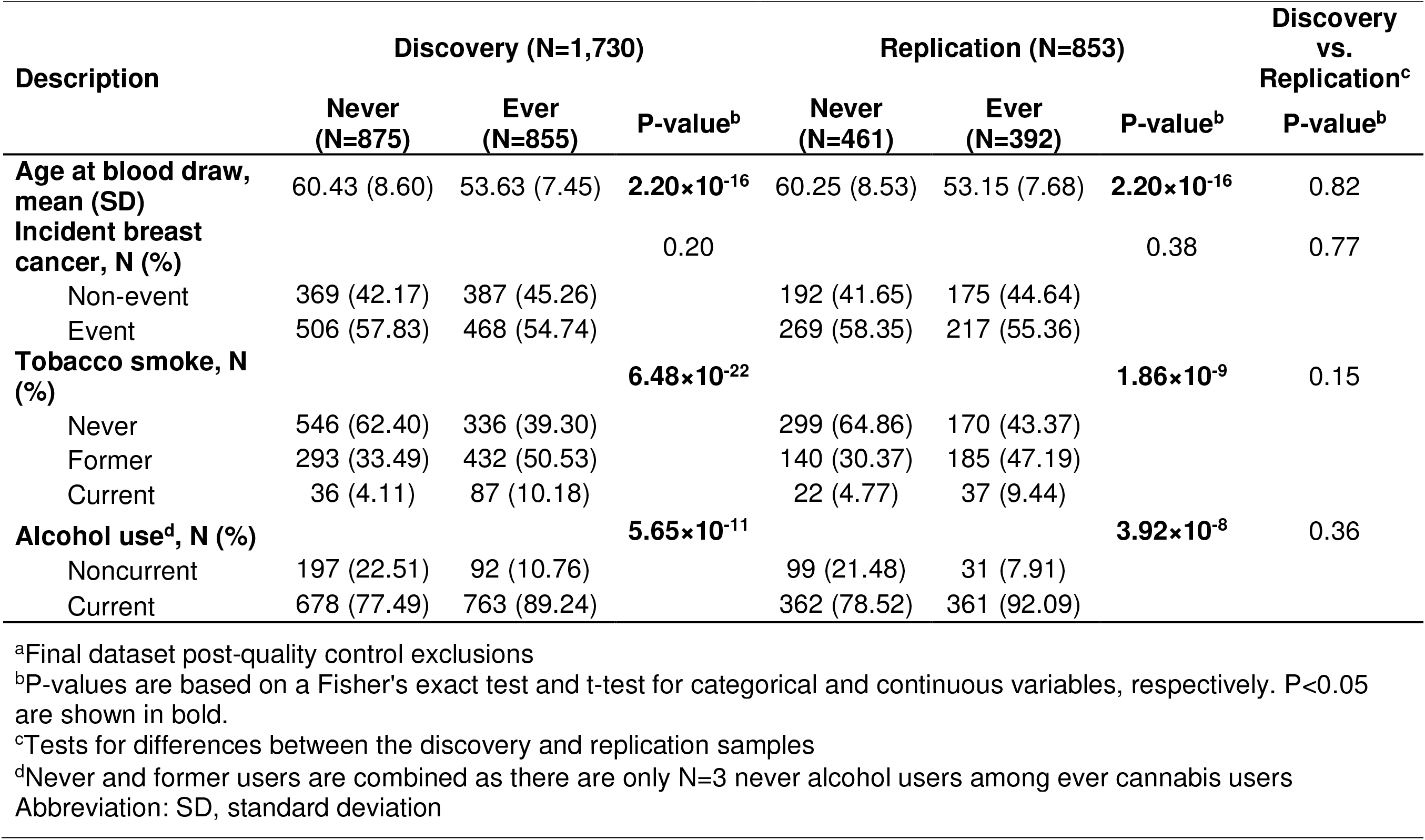
Description of NIEHS Sister Study samples (total N = 2,583)^a^ by ever/never lifetime cannabis use.

For the EWAS, robust linear regression models were implemented using the R package, *MASS^22^*, to test the association between lifetime cannabis use (ever [used at least once in lifetime] vs. never use) and methylation (β-value) at each CpG site, adjusting for age (continuous), incident breast cancer status (event, non-event), tobacco smoking (never, former, current), alcohol use (noncurrent, current), laboratory plate, DNA extraction method, six control SVs, and six blood cell type proportions. To adjust for multiple testing, the false discovery rate (FDR) was controlled at 10% (Benjamini and Hochberg)^23^. Replication analyses were conducted using the same set of covariates. For replication, a Bonferroni correction was applied accounting for the number of CpGs tested.

A series of sensitivity analyses were conducted to assess other possible confounders. We considered the following: current perceived stress (derived from items 2, 6, 7, and 14 from the 14-item Perceived Stress Scale instrument^24^; scale 0–20), family income while growing up (low income, middle income and well-off), body mass index (BMI) as calculated from height and weight measured by examiners at baseline (continuous), and self-reported history of depression (yes, no) (Supplementary Table S1). To further rule out possible effects due to an association with subsequent breast cancer and use of alcohol and/or tobacco, we conducted stratified analyses by these variables using the combined sample for significant EWAS findings. EPISTRUCTURE was implemented using the python toolset, GLINT^25^, to rule out genetic confounding for significant EWAS findings. More specifically, the first four EPISTRUCTURE components (principal components) were computed, while accounting for estimated cell type proportions, and included as covariates in the model to adjust for population stratification.

### Multi-CpG classifier: development and validation

We developed a multi-CpG classifier of lifetime cannabis use within the EWAS discovery sample (N = 1,730) and validated it using the replication sample (N = 853; withheld from model training). Least absolute shrinkage and selection operator (LASSO) regression^26^ was used for model training and variable selection. DNAm β-values were adjusted for the same set of covariates used in the EWAS. Instead of including all CpGs for model training, we selected subsets of CpGs aiming to reduce the amount of noise introduced in the model. As no gold-standard selection criteria exists, two different significance thresholds (P<1×10^−5^ and P<1×10^−4^) from our discovery lifetime cannabis use EWAS were applied to select CpGs for input into LASSO regression. The R package, *glmnet*^27^, was used to implement LASSO regression and 10-fold cross validation for model selection (based on lambda) in the discovery sample. Details regarding the analytical steps and models are provided in Supplementary Methods online. Model performance was evaluated in the discovery sample and independent replication sample, separately, using the R package, *pROC*^28^, to perform a receiver operating characteristic (ROC) analysis. We calculated the 95% confidence interval of the area under the ROC curve (AUC) using 5,000 bootstrap iterations and derived empirical P-values for the AUCs using permutation testing (see Supplementary Methods online)

### Follow-up analyses of replicable EWAS findings

To further characterize replicable EWAS findings, we examined the relationship between DNAm levels and total duration of use among ever users in the combined sample (N = 1,239 [N = 855 from discovery + 392 from replication - 8 with missing information]). Total duration of use was divided into quartiles (upper [> 5 years of use] vs. lower quartile [< 1 year of use]) given the skewed distribution and the fact that less than one year of total use was categorized in the same way (i.e., cannot distinguish between 1 month vs. 6 months of total use). We also evaluated the effect of age of initiation (log transformed) on DNAm levels (N = 1,245 [N = 855 from discovery + 392 from replication - 2 with missing information]; mean +/− SD: 20.85 ± 6.76 years). These secondary evaluations were restricted to these two exposure characteristics given the high degree of missing information (23%) and limited variability for frequency of use among ever users.

Our EWAS findings could be due to underlying genetic effects or the effect of exposure to cannabis. First, to explore whether genetic risk factors for lifetime cannabis use could explain our findings, we used the set of eight genome-wide significant single nucleotide polymorphisms (SNPs) identified in the most recent lifetime cannabis use GWAS meta-analysis (N = 184,765; European ancestry)^15^ and the BIOS QTL browser^29^ to identify CpGs associated with the cannabis use-associated SNPs in blood samples (i.e., determine if lifetime cannabis use-associated SNPs are *cis*-methylation quantitative trait loci [*cis*-meQTLs] in blood). We then examined whether any of the CpGs we identified in our EWAS overlapped with these *cis*-meQTL-CpGs. Of note, *cis*- meQTLs provided through the BIOS QTL browser were identified using data on 3,841 Dutch individuals and modeled without accounting for any phenotype or exposure.

Next, we used the BIOS QTL browser^29^ to identify blood-based *cis*-meQTLs for the lifetime cannabis-use associated CpGs observed in our EWAS. The Sister Study participants have been genotyped using the OncoArray that is customized to optimally capture cancer-associated SNPs^30^, but the 230,000 tagging SNP backbone does not provide sufficient coverage to directly model *cis*-meQTL associations with lifetime cannabis use in the Sister Study case-cohort sample. As such, we again used results from the lifetime cannabis use GWAS meta-analysis (N = 162,082 with 23andMe samples removed)^15^ and performed a look-up of the association between BIOS-implicated *cis*-meQTLs and lifetime cannabis use.

## RESULTS

### EWAS sample characteristics

On average, women who reported lifetime cannabis (ever) use were younger and more likely to report alcohol and tobacco use (Table 1). The prevalence of current tobacco smoking was low (7% of the total sample) and, among ever cannabis users, only half identified as a former tobacco smoker. Alcohol use was more prevalent in the sample with 10% of ever cannabis users reporting noncurrent use and only 3 individuals indicating never use. In addition, ever cannabis users were more likely to grow up in a family with higher income levels, have a lower BMI, and have a higher estimated proportion of CD8 T cells and a lower proportion of NK cells (Supplementary Table S1). However, after controlling for age, the estimated cell type proportions were not associated (P>0.05) with lifetime cannabis use. There was also suggestive evidence in the discovery sample, but not in the replication sample, for differences between ever and never users by self-reported history of depression and current perceived stress (Supplementary Table S1). None of the covariates examined, except for the estimated proportion of NK cells, were significantly different (P>0.05) between the discovery and replication samples (Table 1 and Supplementary Table S1).

### EWAS of lifetime cannabis use: Discovery and Replication

We identified one lifetime cannabis use-associated CpG, cg15973234, at FDR<0.10 (Figure 1 and Supplementary Table S2 [EWAS results with unadjusted P<0.05]). Overall, effects sizes were small (Supplementary Fig. S2), and there was no evidence of inflation (λ=1.02; Supplementary Fig. S3). Further adjustment for potential confounders, including current perceived stress, family income while growing up, BMI, and history of depression did not substantively affect the overall results, thus the more parsimonious model remained our primary model (see Supplementary Fig. S4–S5 for model comparisons).

**Figure 1.**
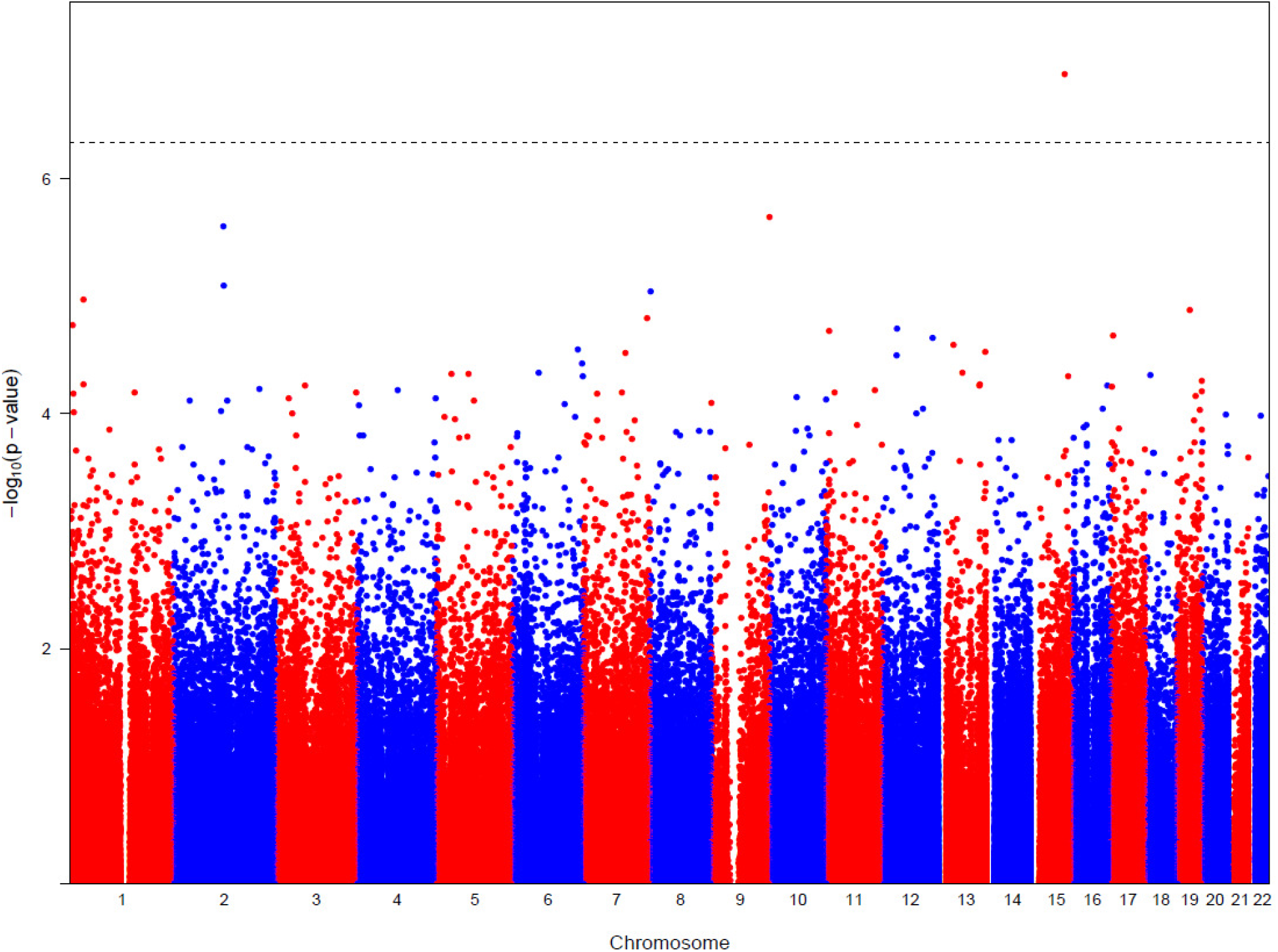
Discovery EWAS of lifetime cannabis use. CpGs are shown according to their position on chromosomes 1–22 (alternating red/blue) and plotted against their -log_10_P- values. The dotted horizontal line indicates FDR<0.10. The genomic inflation factor (λ) was 1.02.

The cg15973234-lifetime cannabis use association met a Bonferroni correction in the replication sample (P_Replication_ = 0.04, β_Replication_ = −0.005; Table 2) and in the combined sample (P_Combined_ = 3.3×10^−8^, β_Combined_ = −0.008; Table 2). In further testing of cg15973234 in the combined sample, we found that it was not associated with total duration of use (P = 0.85; β = 0.0005) or age of initiation (P = 0.35; β = 0.01).

**Table 2.** Replication of cg15973234^a^−lifetime cannabis use association.

| Sister Study Sample | N | Beta | SE | P-value |
| --- | --- | --- | --- | --- |
| Discovery | 1,730 | -0.009 | 0.002 | 1.32×10<sup>-7</sup>,<sup>b</sup> |
| Replication | 853 | -0.005 | 0.003 | 4.51×10<sup>-2</sup> |
| Combined | 2,583 | -0.008 | 0.001 | 3.32×10<sup>-8</sup> |

<sup>a</sup>Probe located within the gene, *CEMIP* (chr15:81072152; GRCh37/hg19 build)
<sup>b</sup>FDR adjusted P-value = 0.06
Abbreviation: SE, standard error

Additional sensitivity analyses designed to further evaluate possible effects of cigarette smoking, alcohol use, incident breast cancer, and genetic ancestry on our observed cg15973234 association did not suggest significant confounding (Supplementary Table S3).

The lifetime cannabis use-associated cg15973234 lies within a CpG island spanning the 5’ untranslated region of the cell migration inducing hyaluronidase 1 (*CEMIP*) gene. The adjacent CpGs in the island are located within 1kb of cg15973234 but are not highly correlated with cg15973234 and thus provide minimal support for the EWAS signal (Supplementary Table S4). The most highly correlated CpG within the region (cg24159335; Pearson correlation [r] with cg15973234 = 0.29) has a similar mean DNAm β-value and is nominally associated with lifetime cannabis use with a direction of effect consistent with cg15973234 (P = 3.2×10^−3^; β = −0.003; Supplementary Table S4).

### Blood-based multi-CpG classifiers of lifetime cannabis use

Using the top 59 most significant CpGs (EWAS discovery P<1×10^−4^) as input into LASSO regression for model training resulted in a classifier composed of 49 CpGs (Supplementary Table S5). We evaluated model performance in the discovery sample used for model training and the independent replication sample. The 49-CpG classifier produced an AUC = 0.74 (95% confidence interval [0.72, 0.76], P = 2.00×10^−5^; Figure 2 and Supplementary Fig. S6) in the discovery sample and AUC = 0.54 ([0.51, 0.57], P = 4.64×10^−2^; Figure 2 and Supplementary Fig. S7) in the replication (validation) sample. Including only the top 5 most significant CpGs (EWAS discovery P<1×10^−5^) for model training resulted in a 3-CpG classifier with reduced model performance (AUC_Discovery_ = 0.59 [0.57, 0.61], P = 2.00×10^−5^; AUC_Replication_ = 0.51 [0.48, 0.54], P = 0.61; Supplementary Table S6 and Supplementary Fig. S8–S10). Our replicable EWAS finding, cg15973234, was included in both models.

**Figure 2.**
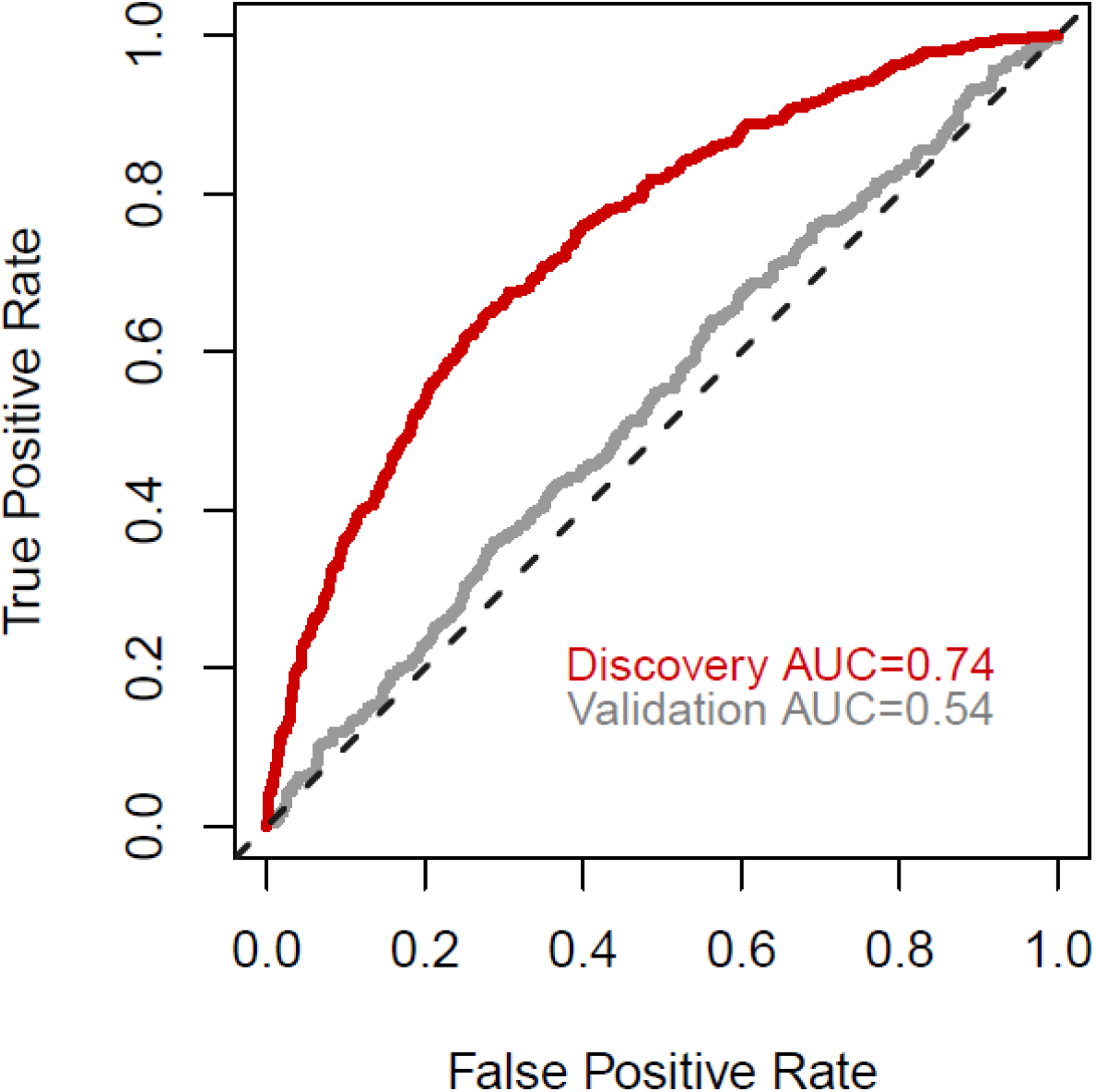
ROC curves for the 49-CpG classifier of lifetime cannabis use in the discovery and replication (validation) samples.

### Follow-up of replicable EWAS finding

None of the eight reported lifetime cannabis use-associated SNPs, from the largest GWAS to date (N = 184,765)^15^, are *cis*-meQTLs of cg15973234 as they are not located on the same chromosome as our EWAS finding. To further assess the possibility that there may be other, weaker genetic risk factors of cannabis use driving our EWAS finding, we used the BIOS QTL browser and identified two independent *cis*-meQTLs for cg15973234 in blood (rs3848177 and rs8025670; FDR<0.05), as determined by statistical modeling of *cis*-meQTL effects^29^. To assess the genetic association between these common SNPs and lifetime cannabis use, we again used results from the lifetime cannabis use GWAS meta-analysis^15^ and found no association (Table 3).

**Table 3.** Independent blood-based *cis*-meQTLs of cg15973234 are not associated with lifetime cannabis use in recent GWAS.

| SNP | Gene | Distance to CpG<sup>a</sup> | *cis*-meQTL P-value<sup>b</sup> | GWAS meta-analysis<sup>c</sup> |  |  |
| --- | --- | --- | --- | --- | --- | --- |
|  |  |  |  | Beta | P-value | MAF |
| rs3848177 | *CEMIP* | 0.064 | 2.37×10<sup>-18</sup> | -0.0024 | 0.85 | 0.16 |
| rs8025670 | *ARNT2* | 188.6 | 5.95×10<sup>-6</sup> | -0.0098 | 0.41 | 0.18 |

<sup>a</sup>Distance in kilobases
<sup>b</sup>N = 3,841; BIOS QTL browser (https://genenetwork.nl/biosqtlbrowser/)
<sup>c</sup>Results based on the International Cannabis Consortium (ICC) + UK-Biobank (UKB) samples (N = 162,082). These SNPs were not among the top 10K results based on the full sample (N = 184,765; ICC + UK Biobank + 23andMe) (Pasman et al. *Nat Neurosci* 2018)
Abbreviation: MAF, minor allele frequency

## DISCUSSION

We report the first blood-based EWAS of cannabis use where we identified and replicated a difference between ever and never users in DNAm levels at cg15973234 (*CEMIP*). DNAm at this site is not correlated with reported cannabis-associated SNPs^15^, suggesting that our finding is unlikely to be genetically-driven and more likely related to the cannabis exposure. Further characterization of the cg15973234-lifetime cannabis use association indicated that the finding appears robust to total duration of use and age of initiation, suggesting that this CpG within *CEMIP* acts as a general indicator of lifetime cannabis use (i.e., a marker of ever vs. never cannabis use that does not vary by duration of use or age at first use).

CEMIP plays a role in hyaluronan binding and degradation^31^. Hyaluronan, one of the main components of the extracellular matrix, is an important regulator of inflammation^32^ and immune processes^33^. Although cannabinoids are thought to play a role in immunoregulation and have anti-inflammatory properties^34^, there is little evidence to date that those processes are modulated by CEMIP or downstream genes in the CEMIP pathway^35,36^. *CEMIP* has been associated with various disorders, including nonsyndromic hearing loss^37^, autoimmune disorders^31,38^, cancer^39–41^, and psychiatric disorders^42,43^. In particular, *CEMIP* has been implicated in bipolar disorder and schizophrenia^43^, as a candidate pituitary gland biomarker for schizophrenia^42^, and as differentially expressed in the striatal tissue of a schizophrenia mouse model (Brd1^+/−^)^44^. These associations are in line with prior reports describing both epidemiological^45,46^ and genetic associations^15^ between cannabis use and psychiatric disorders.

Our work reports the first multi-CpG classifier of cannabis use. Although the 49-CpG classifier produced a statistically significant AUC (empirical P<0.05) in both the discovery (used for model training) and replication (withheld from model training, used for validation) samples, it demonstrated limited discrimination between ever vs. never users in the replication sample (AUC_Replication_=0.54, as compared to AUC_Discovery_=0.74). While a reduction in model performance between training and validation samples is generally expected, these results are also consistent with single CpG association results in our discovery vs. replication samples for top EWAS findings (Supplementary Table S2). None of the CpGs used for model training in the discovery sample were associated (P<0.05) with lifetime cannabis use in the replication sample, apart from the *CEMIP* CpG and one CpG in *SIK3* (SIK family kinase 3). Restricting model training to only the top 5 CpGs based on the discovery EWAS resulted in a 3-CpG classifier with poorer model performance. This pattern of reduced model performance with a smaller set of CpGs has been reported previously, for example, with DNAm biomarkers of alcohol use^47^. While efforts to develop multi-CpG classifiers of alcohol use have been successful (AUC = 0.90–0.99 for the full model [CpGs plus age, sex, and BMI] as compared to AUC = 0.63–0.80 for the null model [age, sex, and BMI]), these classifiers were generated using larger training (N = 2,427) and validation samples (N = 920– 2,003) and focused on an extreme phenotype (current heavy alcohol intake vs. non-drinkers) likely to have much larger effect sizes^47^.

Our study presents the first replicable blood-based CpG biomarker associated with lifetime cannabis use. However, our study has limitations. Our findings may not be fully generalizable due to the case-cohort design (i.e., restricted to non-Hispanic Caucasian women and further enriched for women who subsequently developed breast cancer). Although our study relied on self-reported lifetime cannabis use, which could lead to underreporting and misclassification, these factors would tend to bias towards the null resulting in loss of statistical power. While self-reported age at first use, duration of use, and frequency of use were also collected, we had reduced statistical power (as compared to lifetime use) to evaluate DNAm levels at cg15973234 by duration of use and age of initiation and we were unable to evaluate dose-response due to the low level of response to frequency of use. Further, inaccurate and incomplete recall could affect those data, as the average time since first use was 32.6 ± 6.1 years ago with an average total duration of use of 4.6 ± 7.2 years. Time since last use was not collected to assess recency of use. Finally, we cannot rule out the effect of other illicit drugs (e.g., cocaine, opioids) as those data were not collected in the Sister Study. However, we expect the prevalence of other illicit drug use to be low in this cohort. We also do not suspect that our findings are driven by breast cancer, or the use of tobacco and alcohol because our results were robust to stratified analyses. Cg15973234 was not reported as one of the 2,098 replicated breast cancer-associated CpGs in the most recent breast cancer EWAS^48^. Further, in the largest EWAS of tobacco smoking to date^7^, cg15973234 was not identified as one of the 18,760 CpGs associated with current and/or former smoking (FDR<0.05), or in the largest EWAS of alcohol to date^47^, as one of the 363 CpGs associated with alcohol consumption (P<1×10^−7^).

We are also unable to use our results to make any inference regarding the neurobiological mechanisms underlying cannabis use. In the case of epigenetic biomarkers for substance use there is no *a priori* expectation that genes dysregulated in blood will reflect neurobiological mechanisms underlying substance use. DNAm profiles differ across tissues and cell types and studies conducted using a more biologically relevant brain tissue (e.g., nucleus accumbens, prefrontal cortex) would be needed to inform the neurobiology of cannabis use disorder^49^. Once brain-based DNAm marks in humans (necessitating postmortem tissues) have been identified, however, these results could be used to identify blood proxies of exposure for overlapping blood- and brain-based DNAm changes related to lifetime cannabis use.

Our EWAS results provide evidence that blood-based DNAm can inform cannabis use histories. Larger sample sizes and richer phenotype data (e.g., enabling us to compare more extreme groups, such as lifetime ‘regular’ users) are needed to identify additional CpG biomarkers and to develop stronger, more precise multi-CpG classifiers, including ones indicative of cannabis-related health outcomes.

## ACKNOWLEDGEMENTS

This study was supported by the Intramural Research Program of the NIH, National Institute of Environmental Health Sciences (ZO1 ES-044005 [to DPS] and ZO1 ES-049033 and ZO1 ES-049032 to JAT), as well as the Fellow Program at RTI International (EOJ). These results have been presented previously at the National Institute of Drug Abuse Genetics Consortium meeting held on Jan 14–15, 2019 and posted on bioRxiv. We would like to thank Drs. Alexandra White and Kelly Ferguson for their critical review of the manuscript, and Dr. Grier Page for providing statistical advice.

## DATA AVAILABILITY

De-identified data will be made available by request through the Sister Study Tracking and Review System (https://www.sisterstudystars.org/Default.aspx?projectid=50548533-6eba-4470-83c8-d9019d3a14ad).

