## Supplementary Material for "Epigenome-wide analysis uncovers a blood-based DNA methylation biomarker of lifetime cannabis use"

[1]Center for Omics Discovery and Epidemiology, RTI International, Research Triangle Park, NC, USA; [2]Epidemiology Branch, National Institute of Environmental Health Sciences, Research Triangle Park, NC, USA; [3]Fellow Program, RTI International, Research Triangle Park, NC, USA; [4]Epigenetic and Stem Cell Biology Laboratory, National Institute of Environmental Health Sciences, NIH, Research Triangle Park, NC, USA

*Corresponding author
†Contributed equally

### SUPPLEMENTARY METHODS

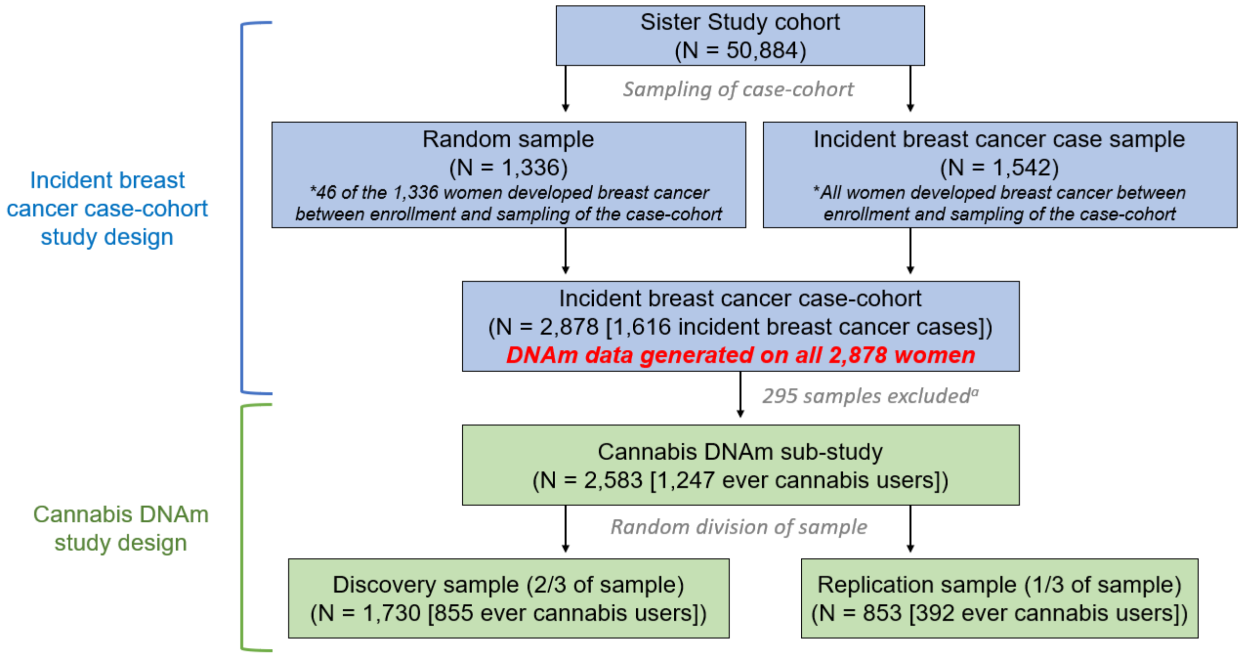


**Supplementary Figure S1.** Sampling strategy used to develop the case-cohort study of 2,878 non-Hispanic white women designed to identify blood-based DNAm changes associated with incident breast cancer (shown above in blue and described previously[1,2]). Design of the cannabis DNAm sub-study is shown above in green. [a]295 samples were excluded due to poor bisulfite conversion efficiency, outlier based on QC diagnostic plots, low call rate, related individuals, missing phenotype data, or date of breast cancer diagnosis preceding blood draw.

**Supplementary Table S1.** Description of NIEHS Sister Study samples (total N=2,583[a]) by ever/never lifetime cannabis use for variables used in sensitivity analyses (income, depression, stress, BMI) or primary model (cell type proportions).

| Description | Discovery (N=1,730) | | | Replication (N=853) | | | Discovery vs. Replication[c] |
|---|---|---|---|---|---|---|---|
| | Never (N=875) | Ever (N=855) | P-value[b] | Never (N=461) | Ever (N=392) | P-value[b] | P-value[b] |
| **Family income level growing up, N (%)** | | | **$1.49\times10^{-7}$** | | | **$6.54\times10^{-6}$** | 1 |
| Poor/low income | 337 (38.51) | 228 (26.67) | | 181 (39.26) | 97 (24.74) | | |
| Middle income/well off | 538 (61.49) | 627 (73.33) | | 280 (60.74) | 295 (75.26) | | |
| **History of depression, N (%)** | | | **$1.12\times10^{-4}$** | | | 0.19 | 1 |
| Yes | 149 (17.03) | 210 (24.56) | | 88 (19.09) | 89 (22.70) | | |
| No | 726 (82.97) | 645 (75.44) | | 373 (80.91) | 303 (77.30) | | |
| **Perceived stress,[d] mean (SD)** | 2.45 (2.68) | 2.71 (2.64) | **$3.75\times10^{-2}$** | 2.52 (2.70) | 2.64 (2.70) | 0.54 | 0.98 |
| **Body mass index, mean (SD)** | 27.93 (6.10) | 27.25 (5.81) | **$1.77\times10^{-2}$** | 28.24 (6.30) | 27.19 (6.03) | **$1.35\times10^{-2}$** | 0.53 |
| **Cell type proportions, mean (SD)** | | | | | | | |
| CD8 T cells | 0.017 (0.027) | 0.020 (0.028) | **$3.34\times10^{-2}$** | 0.016 (0.025) | 0.020 (0.030) | **$2.50\times10^{-2}$** | 0.55 |
| CD4 T cells | 0.175 (0.062) | 0.180 (0.063) | $9.63\times10^{-2}$ | 0.173 (0.059) | 0.179 (0.059) | 0.19 | 0.63 |
| Natural killer cells | 0.064 (0.046) | 0.055 (0.039) | **$1.23\times10^{-5}$** | 0.058 (0.043) | 0.050 (0.038) | **$2.34\times10^{-3}$** | **$5.81\times10^{-3}$[e]** |
| B cells | 0.038 (0.027) | 0.037 (0.027) | 0.63 | 0.038 (0.028) | 0.036 (0.025) | 0.43 | 0.90 |
| Monocytes | 0.047 (0.026) | 0.045 (0.026) | $8.17\times10^{-2}$ | 0.048 (0.028) | 0.046 (0.026) | 0.37 | 0.34 |

| | | | | | | | |
|---|---|---|---|---|---|---|---|
| Granulocytes | 0.614 (0.078) | 0.617 (0.078) | 0.45 | 0.621 (0.078) | 0.621 (0.075) | 0.98 | 0.06 |

[a]Final dataset post-quality control exclusions (N = 295)
[b]P-values are based on a Fisher's exact test and t-test for categorical and continuous variables, respectively. $P<0.05$ are shown in bold.
[c]Tests for differences between the discovery and replication samples
[d]Derived from items 2, 6, 7, and 14 from the 14-item Perceived Stress Scale instrument[3]
[e]$Mean_{Discovery}$ = 0.059 (0.043); $Mean_{Replication}$ = 0.055 (0.041)
Frequencies and means (standard deviations) are presented for categorical and continuous variables, respectively.
Abbreviation: SD, standard deviation

### Multi-CpG classifier development and validation

### Analysis Steps:

1. **Covariate-adjusted DNAm:** We ran the following linear model **(Equation 1)**, separately, for each of 59 CpGs (EWAS P<1×10$^{-4}$) and obtained the residuals, denoted here as $CpG_{resid_i}$ (corresponds to DNAm values adjusted for the same set of biological and technical covariates used in the primary EWAS model [further adjustment for potential confounders, including current perceived stress, family income while growing up, BMI, and history of depression did not substantively affect the EWAS results, thus the more parsimonious model remained our primary model for EWAS and for multi-CpG classifier development]). Models were run separately in the discovery (N = 1,730) and replication (N = 853) samples.

$$CpG = \beta_0 + \beta_1 Age + \beta_2 BreastCancer + \beta_{3-4} Smoking + \beta_5 Alcohol + \beta_6 DNAExtraction + \beta_{7-38} LabPlate + \beta_{39-44} SV + \beta_{45} CD8T + \beta_{46} CD4T + \beta_{47} NK + \beta_{48} Bcell + \beta_{49} Mono + \beta_{50} Gran \quad \textbf{(1)}$$

   $CpG$ corresponds to DNAm β-values, $Age$ is age at blood draw (continuous), $BreastCancer$ is incident breast cancer status (event, non-event), $Smoking$ refers to tobacco smoking (never, former, current), $Alcohol$ is alcohol use (noncurrent, current), $LabPlate$ corresponds to 33 laboratory plates, $SV$ refers to 6 surrogate variables (continuous) of the array control probes, and $CD8T$, $CD4T$, $NK$ (natural killer cells), $Bcell, Mono$ (monocytes), and $Gran$ (granulocytes) correspond to estimated blood cell type proportions (continuous).

2. **Model training in discovery sample (N = 1,730):** Two separate LASSO analyses were run using the discovery sample for model training (fit a logistic regression model for the log-odds):
   a. All 59 $CpG_{resid}$ (EWAS P<1×10$^{-4}$) were included in the LASSO analysis **(Equation 2)**,
   b. Only the top 5 $CpG_{resid}$ (EWAS P<1×10$^{-5}$) were included in the LASSO analysis **(Equation 3)**.

   The R package, *glmnet*[4], was used to implement LASSO regression and 10-fold cross validation for model selection based on lambda. Lambda was selected by optimizing the area under the receiver operating characteristic (ROC) curve (AUC), within 1 standard error of the maximum, in the discovery sample.

$$logit(P_{Cannabis}) = \alpha + \sum_{n=1}^{59} \beta_i CpG_{resid_i} \quad \textbf{(2)}$$

$$logit(P_{Cannabis}) = \alpha + \sum_{n=1}^{5} \beta_i CpG_{resid_i} \quad \textbf{(3)}$$

**3. Evaluation of model performance:**

a. We evaluated model performance in the discovery sample (used for model training) and the independent replication sample (used for validation; withheld from model training), for each model separately

   i. Models:

$$logit(P_{Cannabis}) = \alpha + \sum_{n=1}^{49} \beta_i CpG_{resid_i} \quad \textbf{(4)}$$

$$logit(P_{Cannabis}) = \alpha + \sum_{n=1}^{3} \beta_i CpG_{resid_i} \quad \textbf{(5)}$$

   where regression coefficients for each $CpG_{resid}$ were determined from LASSO (see Supplementary Tables S5 and S6 for coefficients). CpGs with a coefficient of zero are not included in the above counts for Equations 4 and 5 (e.g., 49 CpGs had nonzero coefficients in Equation 4).

b. Model performance was evaluated using the R package, pROC[5], to perform a ROC analysis and calculate the AUC for classifying lifetime cannabis users (function *roc* with default options). We calculated the 95% confidence interval of AUC using 5,000 bootstrap iterations (function *ci.auc* with 'conf.level=0.90, method="bootstrap", boot.n=5000').

c. We conducted permutation testing to derive an empirical P-value for the observed AUCs, following the steps below:

   i. Discovery sample:

      1. Retrieved the ever vs. never cannabis use predictions from the trained models (for the 49-CpG [**Equation 4**] and 3-CpG [**Equation 5**] classifiers, separately)
      2. Permuted cannabis ever/never use status
      3. Calculated AUCs by comparing the predictions from step 1 to the permuted ever vs. never cannabis use outcome
      4. Repeated steps 2–3 50,000 times to obtain an AUC distribution under the null hypothesis of no association between DNAm and cannabis use
      5. Calculated the empirical P-value as the proportion of times we met or exceeded the observed AUC (Supplementary Fig. S6–S7)

   ii. Replication sample:

1. Retrieved the ever vs. never cannabis use predictions for the trained models (for the 49-CpG and 3-CpG classifiers, separately)
2. Permuted cannabis ever/never use status
3. Calculated AUCs by comparing the predictions from step 1 to the permuted ever vs. never cannabis use outcome
4. Repeated steps 2–3 50,000 times to obtain an AUC distribution under the null hypothesis of no association between DNAm and cannabis use
5. Calculated the empirical P-value as the proportion of times we met or exceeded the observed AUC (Supplementary Fig. S9–S10)

### SUPPLEMENTARY RESULTS

**Supplementary Table S2.** Discovery EWAS findings (unadjusted P-value < 0.05) with replication and combined results. **Uploaded separately due to size*

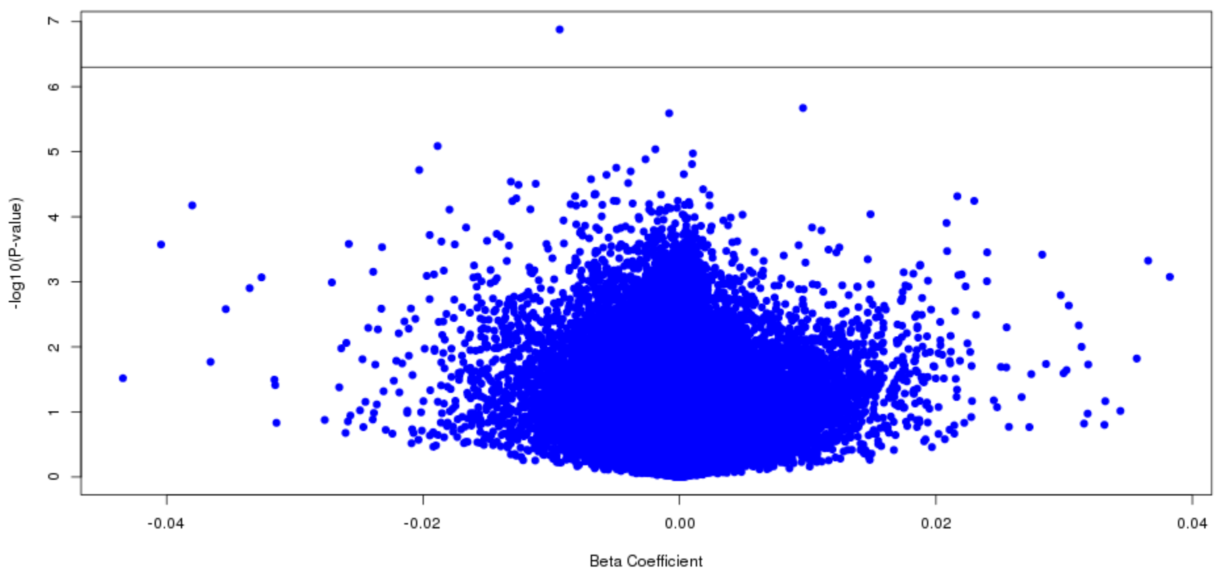


**Supplementary Figure S2.** Lifetime cannabis use EWAS volcano plot. The solid horizontal line indicates genome-wide significance based on FDR<0.10.

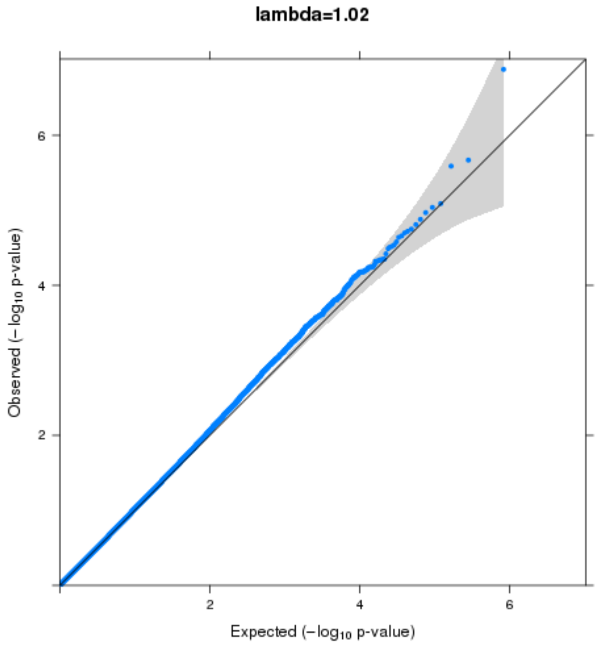


**Supplementary Figure S3.** Quantile-quantile (Q-Q) plot of the lifetime cannabis use EWAS.

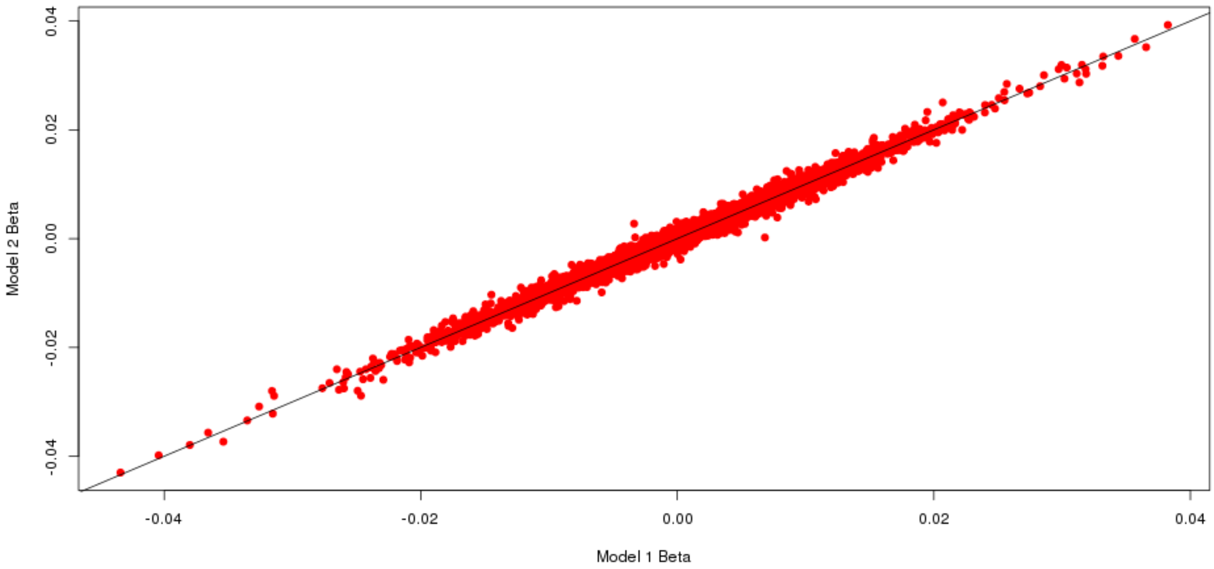


**Supplementary Figure S4.** Sister Study discovery sample: comparison of the lifetime cannabis use coefficient in Model 1 (primary model described in manuscript: DNAm ~ lifetime cannabis use + age + incident breast cancer status + tobacco smoking + alcohol use + laboratory plate + DNA extraction method + 6 negative control SVs + 6 blood cell type proportions) versus Model 2 (further adjustment for potential confounders: current perceived stress, family income while growing, BMI, and history of depression) (Pearson correlation, r = 0.99).

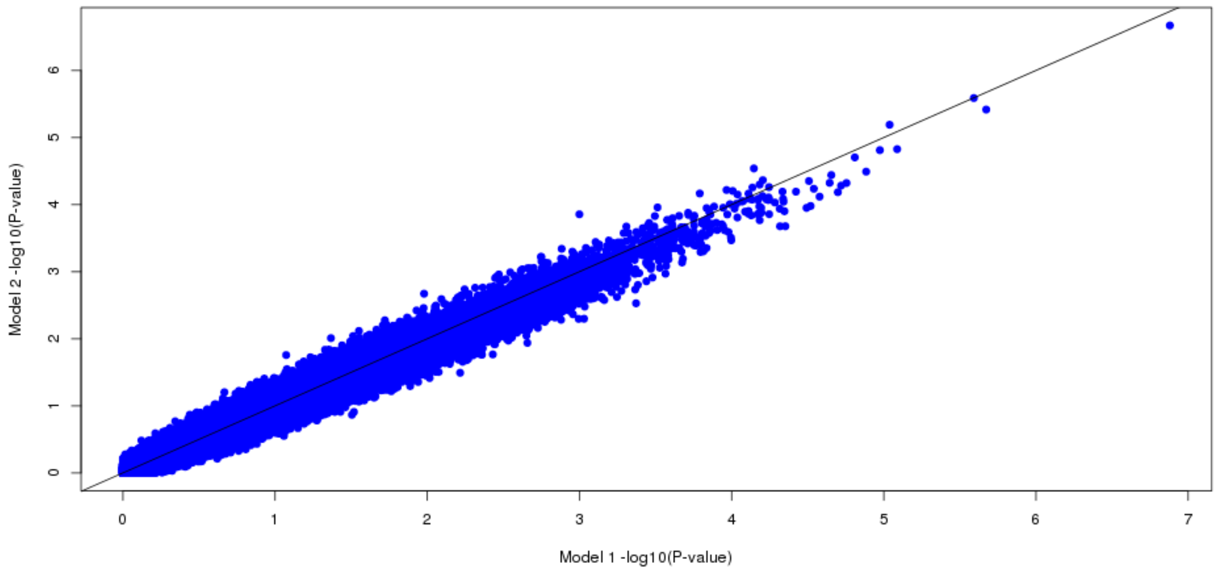


**Supplementary Figure S5.** Sister Study discovery sample: comparison of the lifetime cannabis use $-\log_{10}$(P-value) in Model 1 (primary model described in manuscript: DNAm ~ lifetime cannabis use + age + incident breast cancer status + tobacco smoking + alcohol use + laboratory plate + DNA extraction method + 6 negative control SVs + 6 blood cell type proportions) versus Model 2 (further adjustment for potential confounders: current perceived stress, family income while growing, BMI, and history of depression) (Pearson correlation, r = 0.98).

**Supplementary Table S3.** cg15973234-lifetime cannabis use sensitivity analyses in combined sample.

| Model[a] | Model description | Total N | N ever cannabis users | Beta | SE | P-value[b] |
|---|---|---|---|---|---|---|
| S1 | Additional adjustment for current perceived stress + family income while growing up + body mass index + history of depression | 2,583 | 1,247 | -0.008 | 0.001 | **$8.67\times10^{-8}$** |
| S2 | Excluded breast cancer incident cases | 1,123 | 562 | -0.010 | 0.002 | **$3.74\times10^{-6}$** |
| S3 | Excluded noncurrent alcohol drinkers | 2,164 | 1,124 | -0.007 | 0.002 | **$1.38\times10^{-5}$** |

| | | | | | | |
|---|---|---|---|---|---|---|
| S4 | Excluded current alcohol drinkers | 419 | 123 | -0.016 | 0.004 | **9.61×10$^{-5}$** |
| S5 | Excluded ever tobacco smokers | 1,351 | 506 | -0.009 | 0.002 | **5.47×10$^{-6}$** |
| S6 | Additional adjustment for EPISTRUCTURE PCs 1-4 | 2,583 | 1,247 | -0.008 | 0.001 | **3.15×10$^{-8}$** |

[a]All models were adjusted for age, lab plate, DNA extraction method, negative control SVs 1-6, and estimated cell type proportions (CD8T + CD4T + NK + Bcell + Mono + Gran), except for Model S4 where lab plate was not included as a covariate due to a small N=419 resulting in model convergence issues. Where applicable, tobacco use (Models S1–S4, S6), alcohol use (Models S1-S2, S5-S6), and incident breast cancer status (Models S1, S3–S6) were also included as covariates in the model.
[b]P<0.05 are shown in bold
Abbreviation: SE: standard error

**Supplementary Table S4.** Results for CpGs within the same CpG island as cg15973234.

| CpG | Distance to cg15973234[a] | Beta[b] | SE[b] | P-value[b] | cg15973234 correlation[c] | Mean CpG β-value |
|---|---|---|---|---|---|---|
| cg24159335 | –879 | -0.00276 | 0.0009 | **3.21×10$^{-3}$** | 0.29 | 0.12 |
| cg24796627 | –444 | -0.00018 | 0.0001 | 0.21 | 0.06 | 0.01 |
| cg08177359 | –309 | 0.00062 | 0.0007 | 0.39 | 0.17 | 0.09 |
| cg15973234 | NA | -0.00805 | 0.0015 | **3.32×10$^{-8}$** | NA | 0.12 |
| cg00098718 | 448 | -0.00003 | 0.0002 | 0.87 | -0.0002 | 0.01 |
| cg02273903 | 781 | 0.00002 | 0.0001 | 0.78 | 0.03 | 0.01 |

[a]Distance in base pairs
[b]Based on EWAS of lifetime cannabis use conducted in the combined sample. P<0.05 are shown in bold.
[c]Pearson correlation
Abbreviations: SE: standard error, NA: not applicable

**Supplementary Table S5.** CpGs identified in discovery EWAS (P<1×10$^{-4}$) and used in LASSO regression.

| CpG | Chr | Position[a] | Gene | EWAS Beta[b] | EWAS P-value[b] | LASSO coefficient |
|---|---|---|---|---|---|---|
| cg15973234 | 15 | 81072152 | *CEMIP* | -0.0093 | 1.32×10$^{-7}$ | -3.36 |
| cg13551841 | 9 | 140687144 | *EHMT1* | 0.0096 | 2.12×10$^{-6}$ | 0 |
| cg04195527 | 2 | 118846289 | *INSIG2* | -0.0008 | 2.56×10$^{-6}$ | -24.81 |
| cg03765885 | 2 | 119571674 | *intergenic* | -0.0189 | 8.17×10$^{-6}$ | -1.28 |
| cg04685163 | 8 | 1645455 | *DLGAP2* | -0.0019 | 9.16×10$^{-6}$ | -13.94 |
| cg12710152 | 1 | 32716773 | *LCK* | 0.0010 | 1.06×10$^{-5}$ | 19.11 |
| cg16700555 | 19 | 29700860 | *UQCRFS1* | -0.0027 | 1.31×10$^{-5}$ | -4.80 |

| | | | | | | |
|---|---|---|---|---|---|---|
| cg21804814 | 7 | 152600000 | *intergenic* | 0.0010 | $1.55\times10^{-5}$ | 26.80 |
| cg10328583 | 1 | 6551073 | *PLEKHG5* | -0.0049 | $1.76\times10^{-5}$ | -0.97 |
| cg25343246 | 12 | 34756291 | *intergenic* | -0.0203 | $1.91\times10^{-5}$ | -0.73 |
| cg26870460 | 11 | 6947759 | *ZNF215* | -0.0038 | $2.00\times10^{-5}$ | -1.05 |
| cg01035812 | 17 | 4843585 | *SLC25A11;RNF167* | 0.0003 | $2.21\times10^{-5}$ | 52.64 |
| cg01015899 | 12 | 120663812 | *PXN* | -0.0057 | $2.27\times10^{-5}$ | 0 |
| cg12652442 | 13 | 36738072 | *intergenic* | -0.0069 | $2.64\times10^{-5}$ | -1.90 |
| cg17250160 | 6 | 156919811 | *intergenic* | -0.0131 | $2.88\times10^{-5}$ | -0.92 |
| cg23619365 | 13 | 112712009 | *intergenic* | -0.0040 | $3.03\times10^{-5}$ | -1.18 |
| cg13210467 | 7 | 99775443 | *STAG3;GPC2* | -0.0112 | $3.10\times10^{-5}$ | -0.95 |
| cg17024593 | 12 | 34490115 | *intergenic* | -0.0126 | $3.22\times10^{-5}$ | -2.00 |
| cg01198887 | 6 | 166907883 | *RPS6KA2* | 0.0018 | $3.78\times10^{-5}$ | 2.46 |
| cg16587616 | 6 | 62996022 | *KHDRBS2* | -0.0066 | $4.43\times10^{-5}$ | 0 |
| cg18315960 | 13 | 58204028 | *intergenic* | -0.0066 | $4.50\times10^{-5}$ | 0 |
| cg25668922 | 5 | 34656126 | *RAI14* | -0.0014 | $4.56\times10^{-5}$ | -7.28 |
| cg10586870 | 5 | 75722317 | *IQGAP2* | -0.0066 | $4.59\times10^{-5}$ | -0.97 |
| cg19772897 | 18 | 13263188 | *C18orf1* | 0.0023 | $4.63\times10^{-5}$ | 4.79 |
| cg10176110 | 6 | 168841653 | *SMOC2* | -0.0081 | $4.81\times10^{-5}$ | 0 |
| cg17301216 | 15 | 89920348 | *LOC254559* | 0.0217 | $4.83\times10^{-5}$ | 0.90 |
| cg02473540 | 19 | 58570454 | *ZNF135* | -0.0127 | $5.21\times10^{-5}$ | -1.06 |
| cg07033820 | 1 | 32707210 | *MTMR9L* | -0.0051 | $5.67\times10^{-5}$ | -2.02 |
| cg02235741 | 13 | 99853132 | *UBAC2* | -0.0001 | $5.68\times10^{-5}$ | -75.32 |
| cg03457142 | 3 | 71804859 | *EIF4E3* | 0.0230 | $5.69\times10^{-5}$ | 1.24 |
| cg01039752 | 16 | 81439588 | *intergenic* | -0.0049 | $5.71\times10^{-5}$ | -2.71 |
| cg08390696 | 13 | 99405102 | *SLC15A1* | -0.0130 | $5.74\times10^{-5}$ | -1.05 |
| cg27564939 | 17 | 2207243 | *SMG6;SRR* | 0.0007 | $5.92\times10^{-5}$ | 13.14 |
| cg22835724 | 2 | 205125510 | *intergenic* | -0.0037 | $6.23\times10^{-5}$ | -2.20 |
| cg17464820 | 11 | 116838417 | *SIK3* | -0.0075 | $6.27\times10^{-5}$ | -1.77 |
| cg10520887 | 4 | 96470094 | *UNC5C* | -0.0024 | $6.36\times10^{-5}$ | 0 |
| cg16638540 | 19 | 58570468 | *ZNF135* | -0.0085 | $6.39\times10^{-5}$ | 0 |
| cg07178006 | 11 | 20184718 | *intergenic* | -0.0060 | $6.55\times10^{-5}$ | -0.51 |
| cg21998512 | 7 | 92077031 | *GATAD1* | 0.0004 | $6.55\times10^{-5}$ | 16.17 |
| cg17362109 | 3 | 194981274 | *C3orf21* | 0.0008 | $6.58\times10^{-5}$ | 21.19 |
| cg06221963 | 1 | 154839813 | *KCNN3* | -0.0380 | $6.68\times10^{-5}$ | -0.61 |
| cg13113737 | 1 | 8411074 | *intergenic* | 0.0023 | $6.72\times10^{-5}$ | 10.50 |
| cg16903225 | 7 | 32111066 | *PDE1C* | -0.0081 | $6.74\times10^{-5}$ | -0.76 |
| cg25453681 | 19 | 44100691 | *ZNF576;IRGQ* | -0.0009 | $7.14\times10^{-5}$ | -22.97 |
| cg19402405 | 10 | 64576514 | *EGR2* | -0.0002 | $7.30\times10^{-5}$ | -6.84 |
| cg10010780 | 4 | 187629520 | *FAT1* | -0.0009 | $7.34\times10^{-5}$ | -25.90 |
| cg08145617 | 3 | 32858541 | *TRIM71* | -0.0007 | $7.42\times10^{-5}$ | -45.18 |
| cg08688629 | 10 | 134972932 | *KNDC1* | -0.0116 | $7.67\times10^{-5}$ | -0.60 |
| cg25228746 | 2 | 127865379 | *BIN1* | -0.0015 | $7.74\times10^{-5}$ | -4.53 |
| cg07992500 | 2 | 37896583 | *CDC42EP3* | -0.0179 | $7.75\times10^{-5}$ | -1.46 |

| | | | | | | |
|---|---|---|---|---|---|---|
| cg11325970 | 5 | 87979642 | *LOC645323* | -0.0024 | $7.75\times10^{-5}$ | -9.16 |
| cg25061843 | 9 | 1042970 | *intergenic* | -0.0011 | $8.15\times10^{-5}$ | 0 |
| cg00991794 | 6 | 125284212 | *STL* | -0.0022 | $8.39\times10^{-5}$ | 0 |
| cg25609954 | 4 | 3472246 | *DOK7* | -0.0016 | $8.58\times10^{-5}$ | -25.20 |
| cg01472075 | 16 | 69984927 | *CLEC18A* | 0.0149 | $9.14\times10^{-5}$ | 0.67 |
| cg02705374 | 12 | 97301631 | *NEDD1* | 0.0002 | $9.17\times10^{-5}$ | 92.27 |
| cg03113572 | 19 | 54057415 | *ZNF331* | 0.0049 | $9.30\times10^{-5}$ | 0 |
| cg26987911 | 2 | 113522573 | *CKAP2L* | 0.0004 | $9.57\times10^{-5}$ | 91.23 |
| cg01806956 | 1 | 9460830 | *intergenic* | -0.0061 | $9.84\times10^{-5}$ | -0.01 |

[a]Position based on GRCh37/hg19 human assembly

[b]Discovery EWAS

Abbreviation: Chr: chromosome

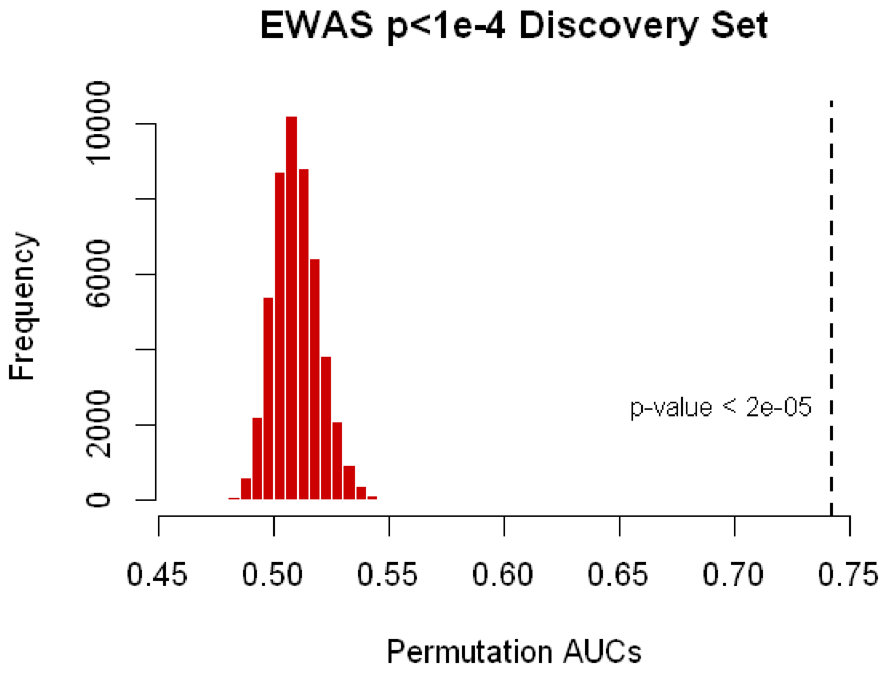


**Supplementary Figure S6.** Permutation testing in the discovery sample (N = 1,730) to obtain a sampling distribution (N = 50,000 permutations) under the null hypothesis. CpGs identified in discovery EWAS ($P<1\times10^{-4}$) were used in LASSO regression. The vertical dotted line indicates the observed AUC.

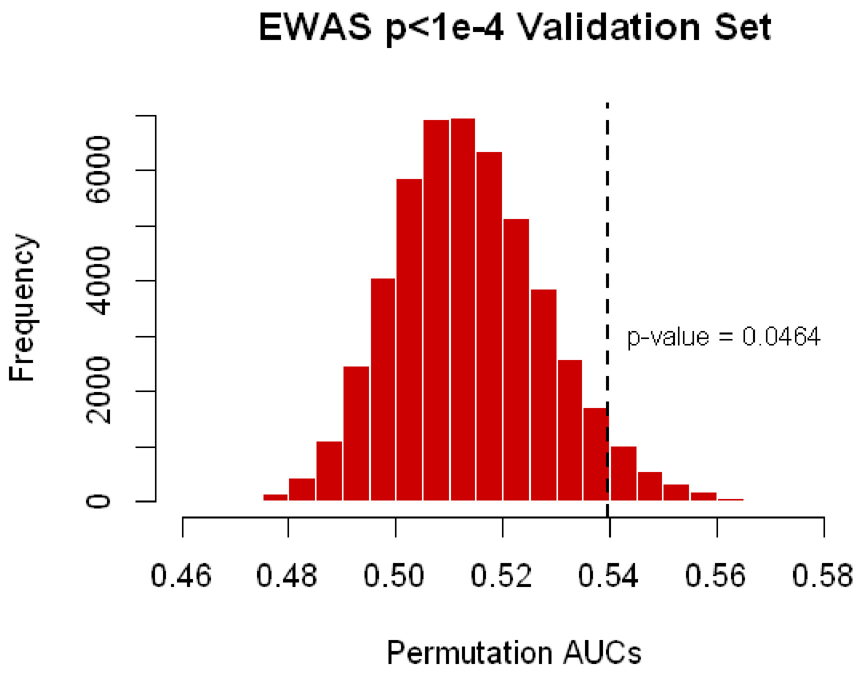


**Supplementary Figure S7.** Permutation testing in the replication (validation) sample (N = 853) to obtain a sampling distribution (N = 50,000 permutations) under the null hypothesis. CpGs identified in discovery EWAS ($P<1\times10^{-4}$) were used in LASSO regression. The vertical dotted line indicates the observed AUC.

**Supplementary Table S6.** CpGs identified in discovery EWAS ($P<1\times10^{-5}$) and used in LASSO regression.

| CpG | Chr | Position[a] | Gene | EWAS Beta[b] | EWAS P-value[b] | LASSO coefficient |
|---|---|---|---|---|---|---|
| cg15973234 | 15 | 81072152 | *CEMIP* | -0.009 | $1.32\times10^{-7}$ | -1.97 |
| cg13551841 | 9 | 140687144 | *EHMT1* | 0.010 | $2.12\times10^{-6}$ | 0 |
| cg04195527 | 2 | 118846289 | *INSIG2* | -0.001 | $2.56\times10^{-6}$ | 0 |
| cg03765885 | 2 | 119571674 | intergenic | -0.019 | $8.17\times10^{-6}$ | -0.82 |
| cg04685163 | 8 | 1645455 | *DLGAP2* | -0.002 | $9.16\times10^{-6}$ | -5.59 |

[a]Position based on GRCh37/hg19 human assembly

[b]Discovery EWAS

Abbreviation: Chr: chromosome

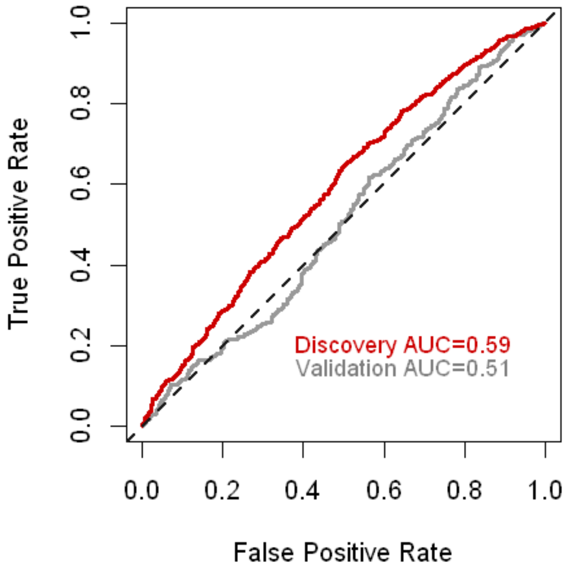


**Supplementary Figure S8.** ROC curves of the 3-CpG classifier of lifetime cannabis use in the discovery and replication (validation) samples.

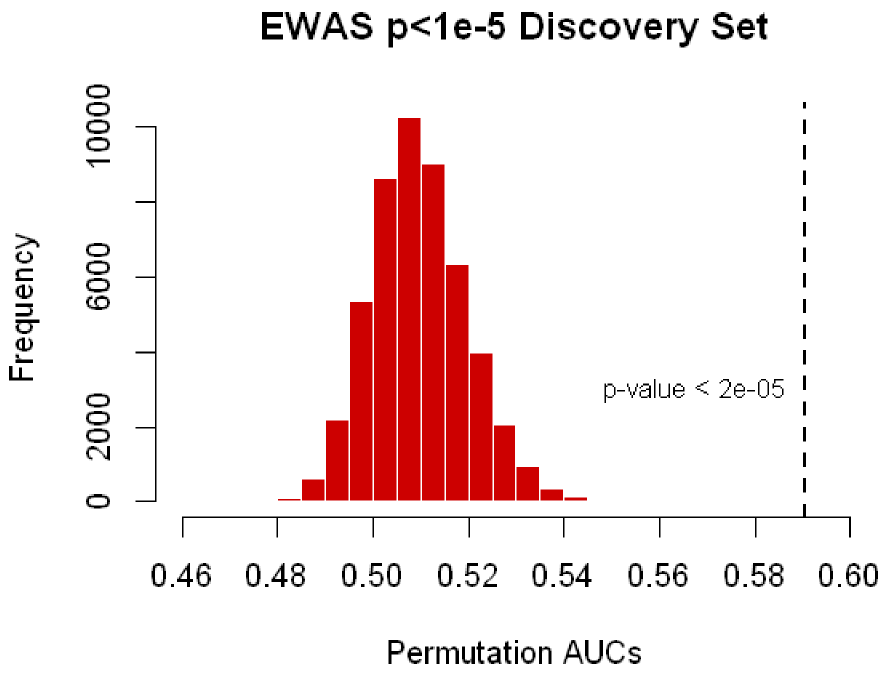


**Supplementary Figure S9.** Permutation testing in the discovery sample (N = 1,730) to obtain a sampling distribution (N = 50,000 permutations) under the null hypothesis. CpGs identified in discovery EWAS ($P<1\times10^{-5}$) were used in LASSO regression. The vertical dotted line indicates the observed AUC.

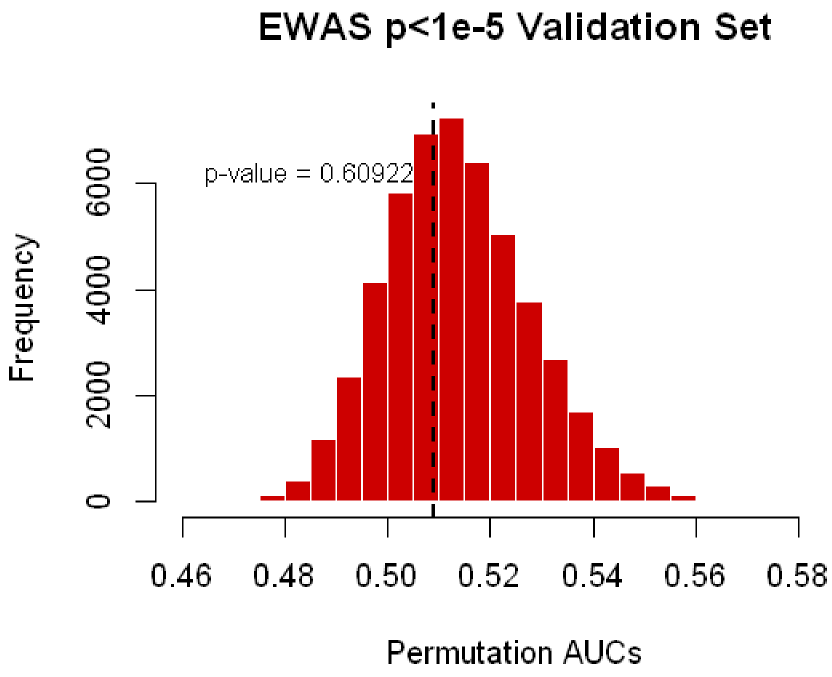


**Supplementary Figure S10.** Permutation testing in the replication (validation) sample (N = 853) to obtain a sampling distribution (N = 50,000 permutations) under the null hypothesis. CpGs identified in discovery EWAS ($P<1\times10^{-5}$) were used in LASSO regression. The vertical dotted line indicates the observed AUC.
